# Annexin A6 modulates the secretion of pro-inflammatory cytokines and exosomes via interaction with SNAP23 in triple negative breast cancer cells

**DOI:** 10.1101/2024.10.22.619710

**Authors:** Nobelle I. Sakwe, Ngoc B. Vuong, Perrin J. Black, Destiny D. Ball, Portia Thomas, Heather K. Beasley, Antentor Hinton, Josiah Ochieng, Amos M. Sakwe

## Abstract

Pro-inflammatory cytokines are secreted via the classical pathway from secretory vesicles or the non-classical pathway via extracellular vesicles (EVs), that together, play critical roles in triple-negative breast cancer (TNBC) progression. Annexin A6 (AnxA6) is a Ca^2+^-dependent membrane-binding protein that in TNBC is implicated in cell growth and invasiveness. AnxA6 is associated with EVs, but whether it affects the secretion of proinflammatory cytokines and/or EVs remains to be fully elucidated. To assess if AnxA6 influences the secretion of cytokines and extracellular vesicles, we used cytokine arrays to analyze secreted factors in cleared culture supernatants from control AnxA6 expressing and AnxA6 downregulated MDA-MB-468 TNBC cells. This revealed the diminished secretion of monocyte chemoattractant protein 1 (MCP-1/CCL2), interleukin 8 (IL-8), dickkopf1 (DKK1), throbospondin-1 (TSP-1), and osteopontin (OPN) following AnxA6 downregulation. We also show that the secretion of small EVs is strongly reduced in AnxA6 downregulated cells and that upregulation of AnxA6 promoted the secretion of treatment was also associated with increased EVs associated Rab7, cholesterol, and MCP-1 levels. Moreover, cholesterol content in EVs was significantly higher in AnxA6-expressing cells than in AnxA6 downregulated cells and following chronic lapatinib induced upregulation of AnxA6. Mechanistically, we demonstrate that the secretion of MCP-1 and/or EVs is AnxA6 dependent and that this requires the translocation of AnxA6 to cellular membranes and its interaction with SNAP23. AnxA6 neutralizing antibodies strongly diminished the survival of AnxA6 low TNBC cells but had minimal effects on the survival of TNBC cells expressing relatively high levels of the protein. Together, these data suggest that AnxA6 facilitates the secretion of EVs and proinflammatory cytokines that may be critical for TNBC progression.

## Introduction

Annexin A6 (AnxA6) is a Ca^2+^-dependent membrane-binding protein that in triple-negative breast cancer (TNBC), is implicated in cell growth (tumor progression), and motility (tumor metastasis). It has also been shown to be involved in calcium (Ca^2+^) and cholesterol homeostasis and drug resistance (1). Interestingly, in TNBC cells, the expression of AnxA6 is relatively higher in mesenchymal-like, pro-invasive cells than in highly proliferative basal-like epithelial TNBC cells (2, 3). We have also shown that the cellular levels of AnxA6 are inducible by Ca^2+^ channel blockers (4), tyrosine kinase inhibitors like lapatinib (5), and by physiological cues like hypoxia (6). Although it lacks a classic signal peptide for secretion, AnxA6 is consistently detected in the extracellular space mostly associated with extracellular vesicles (7, 8) but also as an extracellular vesicle-free protein (9).

Proinflammatory cytokines (PICs) are largely known to be secreted into the tumor microenvironment (TME) by infiltrating immune cells, but are also secreted by tumor cells and other cells in the TME (10). Some of these factors, such as monocyte chemoattractant protein-1 (MCP-1/ CCL2), may promote the recruitment of monocytes, memory T cells, and dendritic cells when secreted by tumor cells (11, 12). Additionally, secretion of interleukin 8 (IL-8/CXCL8), promotes the recruitment of neutrophils to the TME (13). These factors are known to promote the development and progression of breast cancer by influencing inflammation, specific immune responses, cell proliferation, angiogenesis, and metastasis (14, 15). Secretion of cytokines with a signal peptide such as MCP-1 occurs via packaging of the *de novo* synthesized proteins into secretory vesicles or granules, followed by release via receptor-mediated (regulated) or non-regulated (constitutive) exocytosis (16, 17). Cytokines without a signal peptide, on the other hand, are secreted via non-classical exocytosis in which they are packaged into vesicles and released as nano-sized, membrane-bound extracellular vesicles (EVs) (18). While proinflammatory cytokines exert their effects via their receptors, EVs by virtue of their cargo, play an important role in intercellular and cell-extracellular matrix interactions in the TME (19).

AnxA6 containing EVs have been reported to be pro-metastatic and mediate drug resistance by mechanisms that include interaction with cell surface receptors, with membrane phospholipids and either specific extracellular matrix (ECM) components or F-actin, and activation of signaling effectors such as focal adhesion kinase, nuclear factor kappa B (NFkB), and induction of autophagy (20, 21, 1, 9, 22). Like most annexins, AnxA6 translocates to the plasma membrane in a Ca^2+^-dependent manner. Consequently, altered expression of AnxA6 may affect several membrane-associated pathways and cellular processes including vesicle trafficking, exocytosis (5) and cholesterol homeostasis (23).

A recent report suggests that cells secrete exosomes and other EVs upon plasma membrane damage and that AnxA6 depletion stalls MVBs at the cell periphery (24). Interestingly, N- and C-terminal truncated AnxA6 localize to different membranes, suggesting that AnxA6 may tether MVBs to the plasma membrane (24). However, whether it influences the secretion of pro-inflammatory cytokines and/or extracellular vehicles (EVs) independent of cell damage remains to be fully elucidated. In this study, we assessed whether altered expression of AnxA6 by RNA interference and/or chronic lapatinib-induced expression of AnxA6 influences the secretion of proinflammatory cytokines and EVs in TNBC cells. We demonstrate that the secretion of MCP-1 and other proinflammatory cytokines and/or extracellular vesicles is AnxA6 dependent and that this requires the translocation of AnxA6 to cellular membranes and the interaction of AnxA6 with SNAP23. AnxA6 neutralizing antibodies strongly diminished the survival of AnxA6 low TNBC cells but had minimal effects on the survival of TNBC cells expressing relatively high levels of the protein. Together, these data suggest that the secretion of proinflammatory cytokines and EVs, as well as cholesterol enrichment in EVs is AnxA6-dependent, and that its interaction with SNAP23 promotes the secretion of these factors, which may underlie TNBC progression and poor patient outcomes.

## Materials and Methods

### 2.1 Cell Culture

A non-malignant breast cell line, MCF 10A and breast cancer cell lines BT 549, MDA-MB-468, MDA-MB 231, HCC 1806 cells, were all purchased from American Type Culture Collection (ATCC, Manassas, VA). MCF 10A, BT 549, and HCC 1806 were cultured in Dulbecco’s modified Eagle’s nutrient F-12 (DMEM/F12) medium (Gibco, Life technologies, Grand Island, NY), while MDA-MB 231 was grown in RPMI 1640 (Gibco, Life technologies, Grand Island, NY) and MDA-MB-468 in Leibovitz’s L15 (Gibco, Life technologies, Grand Island, NY) media. All media were supplemented with 10% fetal bovine serum, NaHCO3 (10 mM), penicillin (100 units/ml), and streptomycin (50 units/ml) until 70-80% confluent. The cells were maintained in a humidified 95% air and 5% CO_2_ incubator at 37 °C and media were changed every 2–3 days. Where indicated, serum starvation was carried out by culturing the cells in base medium supplemented with 0.5% FBS and the indicated antibiotics. For treatment of cells with the indicated compounds, the cells were trypsinized, seeded at desired cell density and allowed to attach overnight in complete medium. The following day, the media were aspirated and replaced with fresh media containing the indicated concentrations of the drugs and for the indicated times.

### 2.2 Plasmid transfection

The generation of AnxA6 downregulated MDA-468 and BT 549 cells as well as AnxA6 overexpressing MDA-468 and HCC1806 cells were previously described (3, 5, 8). Briefly, cells were transfected with non-silencing control (NSC) and AnxA6 coding sequence targeting shRNAs (A6sh2 and A6sh5) cloned into the pGIPZ lentiviral vector (Horizon Discovery), or the empty vector and flag tagged AnxA6 in plasmid pLV[Exp]-EGFP:T2A:Puro-CMV (Vectorbuilder). Transfected cells were sorted for green fluorescent protein expression and then selected with puromycin for three to five weeks. The stably transfected MDA-468 cells were then treated with lapatinib (2 μM) over several months to generate lapatinib resistant cells (LAP-R) and incubated without lapatinib for five days to generate lapatinib withdrawn cells (LAP-RW) as previously described (5).

### 2.3 Culture Supernatant

Cells were cultured to 80% confluency, they were passaged, and the same number of cells (∼1-2 × 10^6 cells/ml) seeded into T-75 flasks and incubated overnight at 37°C. The following day, cells were washed twice with room temperature PBS and cultured in 10 ml of serum free medium overnight. The culture supernatant was transferred into new tubes, cleared by centrifugation at 275 x g for 5 mins and aliquoted into 2ml tubes for storage at -80°C or used in antibody arrays and isolation of EVs. Then the cells were harvested by trypsinization, and cell count performed using an automated cell counter TC20™ (Bio-Rad Laboratories, Hercules, CA).

### 2.4 Cytokine Array

Cytokine profiling was performed by using the Proteome Profiler Human XL Cytokine Array Kit according to the manufacturer’s instructions (R &D Systems, Inc., Minneapolis, MN, USA, Cat #ARY022B), and cleared culture supernatants collected from AnxA6 expressing (NSC) and AnxA6 depleted (A6sh5) MDA-468 cells. The blots were revealed by enhanced chemiluminescence (ECL) (Perkin Elmer, Waltham, MA), and the intensity of the spots was quantified by using ImageJ.

### 2.5 ELISA Assays

Quantification of Human CCL2/MCP-1 and Human IL-8/CXCL8 concentration in TNBC cell lysates, culture supernatants or isolated EVs, was done using the DuoSet^R^ sandwich ELISA Kits, following the manufacturer’s instructions (R&D Systems, Inc., Minneapolis, MN, USA). The reaction was stopped by addition of 2N H_2_SO_4_ and the absorbance was measured at 450 nm. The concentration of cytokines was extrapolated from a standard curve.

### 2.6 Co-Immunoprecipitation and Western blotting

Cells were cultured to 80% confluency, trypsinized, and washed twice with ice-cold phosphate buffered saline (PBS). Cells were then resuspended in radioimmunoprecipitation assay (RIPA) buffer (50mM Tris-HCl, pH 7.4, 1% NP-40, 0.1% sodium deoxycholate, 150mM NaCl, 1mM EDTA) supplemented with protease inhibitor cocktails (Sigma) and phosphatase inhibitors (20mM sodium fluoride, 50 mM β-glycerophosphate, and 1μM sodium ortho-vanadate) and incubated on ice for 30mins. After centrifugation at >10,000 × g for 10 mins at 4°C, the protein concentration in the whole cell lysate or various subcellular fractions was determined by using the Bradford assay (Bio-Rad Laboratories, Hercules, CA). Immunoprecipitation was performed as previously described (3) and the bound proteins were analyzed by western blotting using the following antibodies: anti-AnxA6 (Santa Cruz Biotechnology sc-271859), anti-β-actin (Sigma, A1978), EGFR (Cell Signaling Technology, C74B9), pEGFR (Cell Signaling Technology, D95F2), GAPDH (Santa Cruz Biotechnology sc-25778), CD63 (Proteintech, 25682-1-AP, USA), Rab7 (Cell Signaling Technology, D7A5), Flotillin-1 (Santa Cruz sc-25506), SNAP23 (Abcam, ab-131242 ER 8538), anti-hCCL2/MCP-1 (R & D Systems, MAB679), anti-hCXCL8/IL-8 (R & D Systems, MAB208). The blots were revealed by (ECL) (Perkin Elmer, Waltham, MA) and the intensity of the bands was quantified by using ImageJ.

### 2.7 GST Pulldown assay

Control and GST (glutathione S-transferase) fusion constructs of AnxA6 were expressed in *E. coli* BL21 and induced with 1 mM isopropyl-1-thio-β-D-galactopyranoside (IPTG) at 30°C for overnight. The cells were harvested by centrifugation at 4500×g for 10 min and the pellets were resuspended in lysis buffer containing lysozyme to a final concentration of 0.5 mg/ml. After incubation for 1 h on ice, the lysates were homogenized using a Dounce Homogenizer (10 strokes). The GST construct was recovered from the supernatant after centrifugation at 12,000×g for 45 min (GST fraction). The supernatants with the GST or GST-AnxA6 fusion protein were incubated with glutathione-Sepharose 4B beads (GE Healthcare Life Sciences) for 2 h at RT, washed 3 times with PBS, and maintained at 4°C. 20 μg of GST or GST-AnxA6 fusion protein, immobilized to glutathione beads, were incubated with 500 μg of total protein from MDA-468 cells (prepared in lysis buffer with protease inhibitors). The beads were washed three times in lysis buffer, and proteins were eluted in SDS sample buffer and analyzed by immunoblotting.

### 2.8 Isolation of small extracellular vesicles (exosomes)

Cells were grown to 70-80% confluency, and then incubated overnight in exosome-free medium. The culture supernatant was centrifuged at 1500 x *g* for 4 min to remove cells and cell debris and filtered through 0.45μ filters. EVs were isolated from the cleared culture supernatants by differential velocity centrifugation as previously described (7,8). The size and concentration of isolated EVs were determined by Nanosight tracking analysis using ZetaView PMX 110 (Particle Metrix, Germany), and the expression of AnxA6 and other exosome markers was assessed by western blotting.

### 2.9 Transmission electron microscopy

The density-gradient-purified small EV in PBS were spotted on Formvar-coated grids (Electron Microscopy Sciences) the grids for 15 min, then fixed with 4% paraformaldehyde for 10 min, and washed with PBS. Free aldehyde groups were quenched with 0.15 M glycine in PBS and the grids were stained with 2% phosphotungstic acid, pH 6.1 and allowed to air-dry. For immunogold staining, the grids were blocked with 5% bovine serum albumin (BSA) in PBS and sequentially incubated with mouse anti-AnxA6 antibody and colloidal gold-conjugated donkey anti-mouse IgG and then stained with 2% phosphotungstic acid. The negative-stained samples were imaged on a FEI Tecnai T12 Transmission Electron Microscope at 100 kV using an AMT CR41 side mounted CCD camera.

### 2.10 Cholesterol measurements

Cholesterol in the isolated EVs was determined by using equal volumes of the isolates, equal numbers of particles and the Amplex™ Red Cholesterol Assay Kit (Life Technologies) according to the manufacturer’s instructions and as previously described (5). The converted fluorescent resorufin was measured by using a fluorescent microplate reader at 560/590 nm (Ex/Em) and the levels of cholesterol were extrapolated from a cholesterol standard curve.

### 2.11 Proximity Ligation Assay

Cells were plated in 8 well chamber slides and cultured overnight in complete medium. In situ PLA for AnxA6 and SNAP23 was conducted by using the Duolink^®^ In Situ Red starter kit Mouse/Rabbit with mouse anti-AnxA6 (Santa Cruz) and rabbit anti-SNAP23 (Abcam) as described by the manufacturer (Sigma-Aldrich). The reactions were visualized by confocal microscopy using a 40X magnification.

### 2.12 Statistical analysis

Experiments were carried out at least three times, and data represented as mean values with standard deviation, except otherwise indicated. ELISA data was analyzed using Microsoft Excel 2016. The results of ELISA and WB were compared by two-tailed unpaired Student’s t-test and the difference was considered significant if p<0.05.

## Results

### 3.1 AnxA6 influences the secretion of pro-inflammatory cytokines and extracellular vesicles in TNBC cells

To demonstrate if AnxA6 mediates cytokine secretion and extracellular vesicles (EVs) in TNBC cells, we first profiled secreted proteins in cleared culture supernatants from control AnxA6 expressing (NSC) and AnxA6 downregulated (A6sh) MDA-468 cells by antibody arrays **(Fig. 1A**). As shown in **Fig. 1B**, reduced expression of AnxA6 inhibited the secretion of DKK-1, IL-8, MCP-1, OPN and TSP-1, but stimulated the secretion of KLK5. Among these factors, the secretion of MCP-1 in MDA-468 cells was abolished following down-regulation of AnxA6. By using breast cancer gene-Expression Miner v3.0 (25), we show that AnxA6 and CCL2 (MCP-1) have a positive Pearson’s pairwise correlation of (0.30, p<0.0001) for all TNBC subtypes (n=699) (**Fig. 1C**). To validate the expression and secretion of MCP-1 in TNBC cells distinct AnxA6 expression levels, we show that MDA-468 that expresses the lowest level of AnxA6 and a prototype basal-like TNBC cell line express the highest level of MCP-1 **(Fig. 1C)**, and that the secretion of MCP-1 is AnxA6 dependent in both BT-549 and MDA-468 TNBC cells **(Fig. 1D)**.

**Figure 1.**
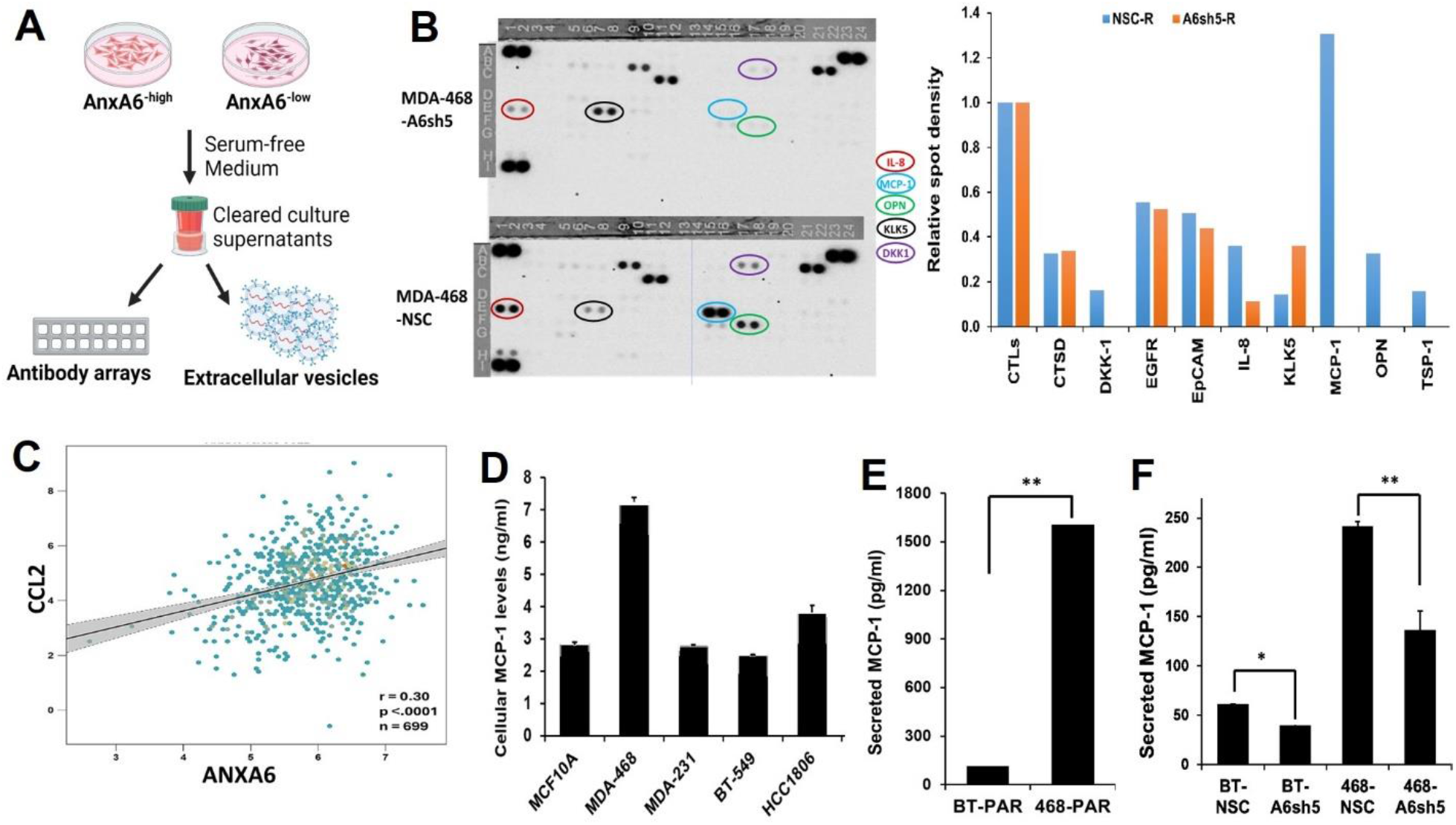
Secretion of pro-inflammatory cytokines is AnxA6 dependent in TNBC cells. **(A)** Schematic showing the experimental design to harvest cleared 24 h culture supernatants from control (NSC) and AnxA6 downregulated (A6sh5) MDA 468 cells. **(B)** Detection of secreted proteins by antibody arrays. Quantification of secreted proteins by ImageJ particle analysis. Bars represent the spot density relative to controls (CTLs). **(C)** Validation of AnxA6-dependent secretion of MCP-1. Cell type-specific expression of MCP-1 in TNBC cells. **(D)** Secreted MCP-1 in control (NSC) and AnxA6 downregulated (A6sh5) BT-549 and MDA-468 cells.

To determine if AnxA6 expression status also influences the secretion of EVs, we isolated EVs from cleared culture supernatants of control AnxA6 expressing, and AnxA6 downregulated BT-549 and MDA-468 TNBC cells by differential velocity centrifugation. **Fig. 2A** shows the transmission electron micrographs of isolated EVs. Analysis of the isolated EVs by using the ZetaView® Nanoparticle Tracking Analyzer revealed that downregulation of AnxA6 strongly inhibited the secretion of EVs in both BT-549 and MDA-468 cells **(Fig. 2B and C**). Upregulation of AnxA6 in the AnxA6-low MDA-468 cells led to increased secretion of EVs (**Fig. 2D**). Quantification of cholesterol in the isolated EVs revealed that downregulation of AnxA6 in these cell lines is associated with decreased EV-associated cholesterol **(Fig. 2E)**, while upregulation of AnxA6 in MDA-468 cells led to increased cholesterol in EVs **(Fig. 2F)**. Together, this suggests that like MCP-1 and other proinflammatory cytokines, the secretion of EVs and cholesterol loading into EVs is AnxA6-dependent in TNBC cells.

**Figure 2:**
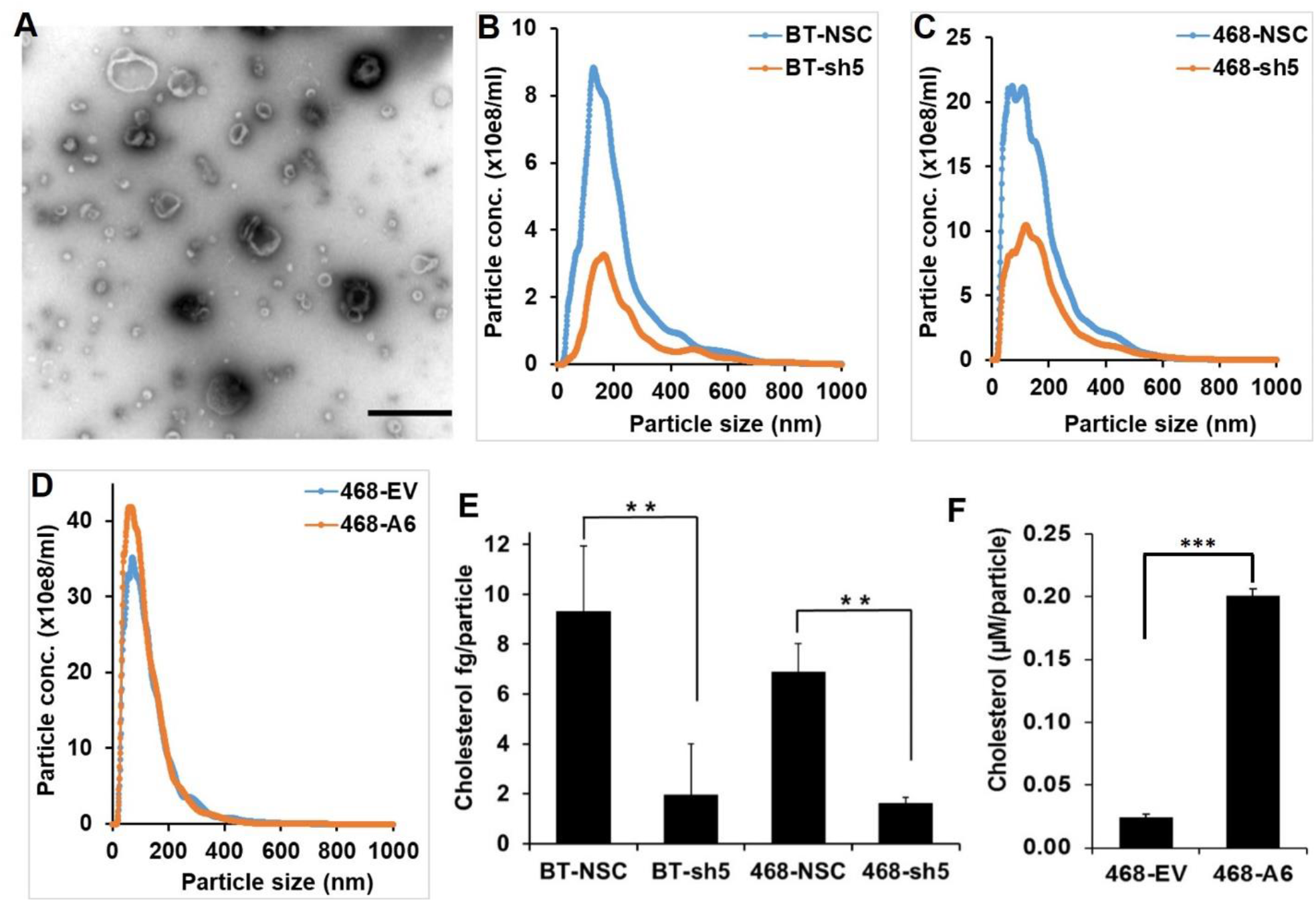
Secretion of small extracellular vesicles is AnxA6 dependent in TNBC. The altered expression of AnxA6 affects the secretion and enrichment of cholesterol in EVs from TNBC cells. Extracellular vesicles (EVs) were purified from culture supernatants from control (NSC) and AnxA6 downregulated (sh5) BT-549 or MDA-468 TNBC cells and from empty vector control and AnxA6 upregulated MDA-468 cells. **(A-C)** EV particle analyses. **(D)** Verification of isolated EVs by transmission electron microscopy. Bars denote 500nm. (E-F) Determination of cholesterol in EVs from control and AnxA6 downregulated TNBC cells (E) and empty vector control and AnxA6 upregulated MDA-468 cells **(F)**. Bars represent EV-associated cholesterol/particle. ** denotes p<0.01

We have previously reported that chronic lapatinib treatment of AnxA6-low MDA-468 TNBC cells is associated with AnxA6 upregulation and accumulation of cholesterol in late endosomes (5). To determine if chronic lapatinib-induced expression of AnxA6 is associated with increased secretion of EVs and proinflammatory cytokines, we first confirmed that lapatinib inhibited EGFR activation and that AnxA6-expression is upregulated in the chronically lapatinib treated cells (**Fig. 3A**). Analysis of the isolated EVs by western blotting revealed that AnxA6 and the small Ras-related GTPase Rab7, were enriched in EVs isolated from lapatinib treated cells (**Fig. 3B**). This analysis also shows that flotillin, an exosome marker was enriched in lapatinib resistant cells while CD63 was strongly down-regulated in AnxA6 depleted and lapatinib resistant cells. (**Fig. 3B**). We also show that EVs-associated cholesterol levels were higher in control AnxA6 expressing cells compared to AnxA6 downregulated cells and more importantly, cholesterol levels in lapatinib treated cells (with increased AnxA6 expression) were significantly higher than in the untreated control cells (**Fig. 3C**). We next show that the secretion of MCP-1 is also AnxA6 dependent following lapatinib treatment and that withdrawal of lapatinib from lapatinib resistant cells, which is accompanied by return of AnxA6 to basal levels and increased cell proliferation (8), was accompanied by significantly higher secretion of MCP-1. (**Fig. 3D**). Assessment of the relative dependency of these genes to AnxA6 revealed that CD63 and AnxA6 have the strongest positive correlation in mesenchymal-like TNBC cells. Together, these observations suggest that MDA-468 cells secrete CD63 enriched EVs while AnxA6 downregulation with or without lapatinib treatment secrete CD63 low EVs.

**Figure 3:**
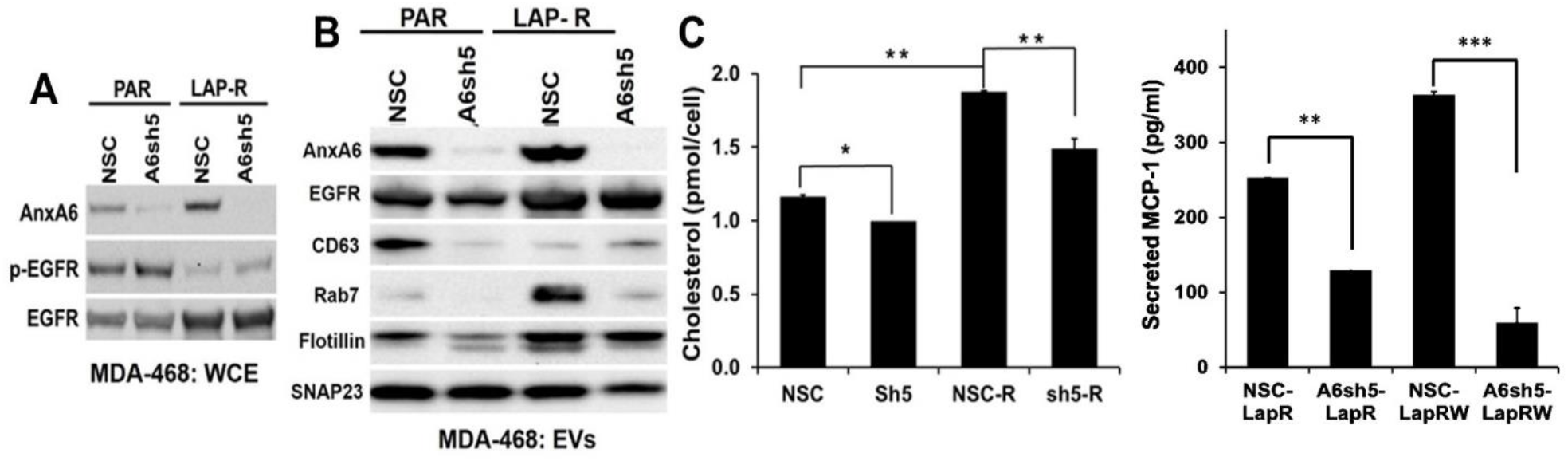
Lapatinib induced expression of AnxA6 is associated with the secretion of AnxA6 and cholesterol enriched EVs. Chronic lapatinib (Lap-R) induced expression of AnxA6 promotes the loading of cholesterol in EVs from MDA-468 cells. **(A)** Western blot showing inhibition of EGFR activation in MDA-468 cells following chronic treatment with lapatinib. **(B)** Enrichment of AnxA6 and RAB7 in EVs from lapatinib resistant MDA-468 cells. **(C)** AnxA6 dependent enrichment of cholesterol in lapatinib resistant MDA-468 cells. Bars represent cholesterol levels/10^6 cells. **(D)** Lapatinib withdrawal leads to increased secretion of MCP-1 in AnxA6 expressing NSC cells (more apparent in NSC-LapRW than the NSC-LapR), and decreased secretion in AnxA6 downregulated A6sh5-LapRW and A6sh5-LapR.

### 3.2 AnxA6 interactome reveals interaction of AnxA6 with the SNARE complex

We used an unbiased proteomic analysis of AnxA6 interactome in which flag-tagged AnxA6 was co-immunoprecipitated with interacting proteins via anti-Flag-M2 antibody (Sigma). Analysis of the interacting proteins revealed that AnxA6 interacted with several proteins and protein families (Table 1). Among these were members of the SNARE (Soluble N-ethylmaleimide-sensitive factor (NSF) Attachment Protein Receptor) membrane fusion complex (26) including SNAP23, VAMP8, and Syt7. We carried out GST Pulldown and Co-immunoprecipitation (Co-IP) and proximity ligation assays to validate the interaction of AnxA6 and SNAP23. As shown in **Fig. 4**, AnxA6 and SNAP23 interacted when immunoprecipitated with either anti-Flag-M2 (**Fig. 4A**) or anti-AnxA6 (**Fig. 4B**) antibodies. A proximity ligation assay using mouse AnxA6 antibody and Rabbit anti-SNAP23 antibody also revealed a strong interaction of AnxA6 and SNAP23 (**Fig. 4C**). Finally, we carried out GST pull-down assays to confirm that AnxA6 interacted with SNAP23 as well as other identified interacting proteins including flotillin, E-cadherin, EGFR and Rab7 (**Fig. 4D**). Together, these data suggest that AnxA6 interacts with at least one component of the SNARE complex with implications in the secretion of proinflammatory cytokines and EVs in TNBC cells

Based on these data, we next used immunogold transmission electron microscopy to determine the localization of AnxA6 in TNBC cells treated with or without Ca^2+^. Consistent with our previous report (9) this analysis revealed that in AnxA6 expressing TNBC cells, extracellular Ca^2+^ treatment induced the translocation of AnxA6 to not only the plasma membrane but also several membrane-bound organelles, including mitochondria, vesicles, and endosomes (**Fig. 5A and B**). As expected, reduced expression of AnxA6 led to its reduced localization to these membrane structures in the cells (**Fig. 5C and D**). Therefore, the translocation of AnxA6 to various membranes supports the relevance of AnxA6 in various membrane-associated processes, including trafficking of intracellular vesicles, uptake of proteins into various organelles, and exocytosis.

**Figure 4:**
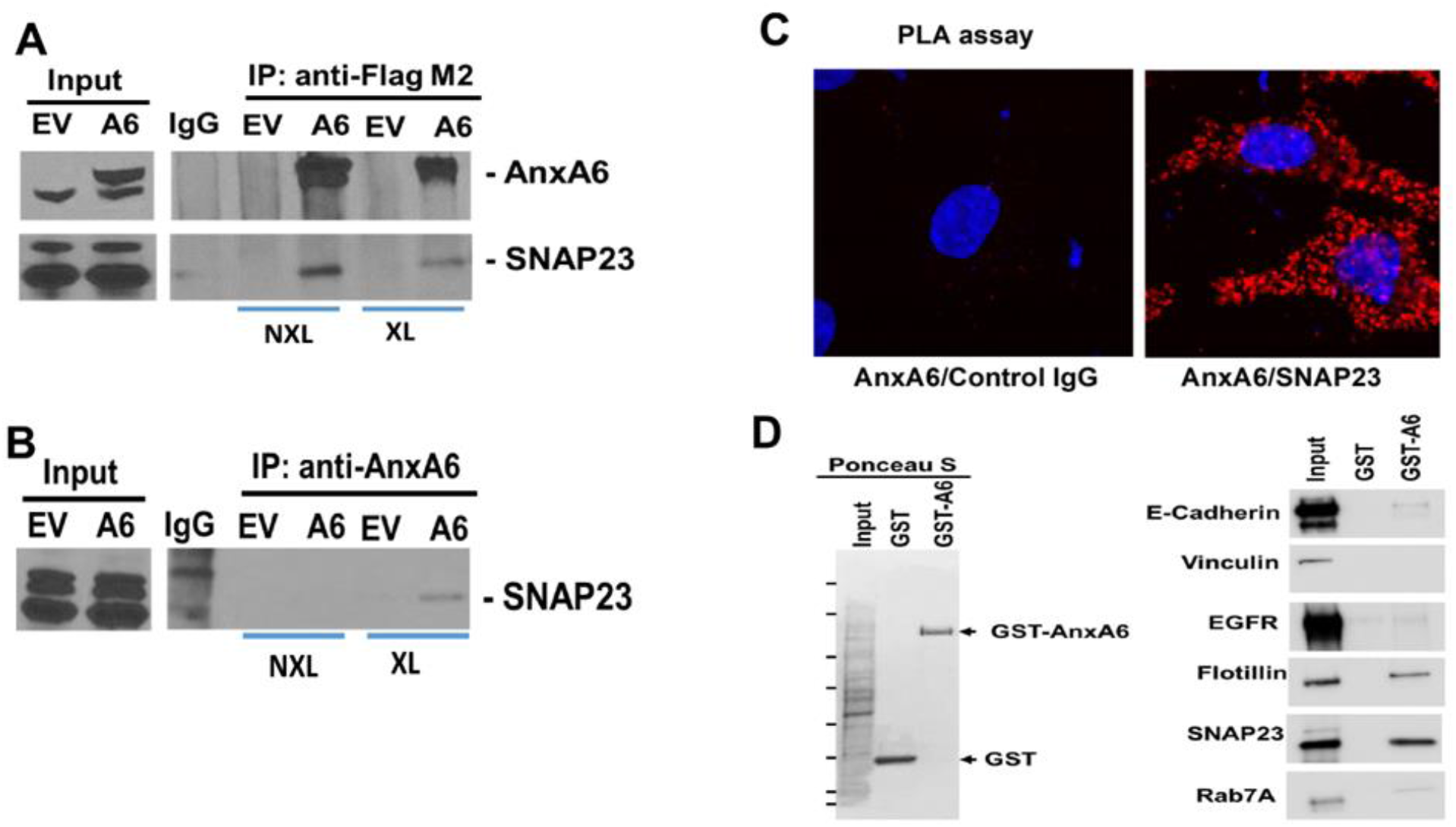
AnxA6 interacts with the SNARE protein SNAP23 in TNBC cells. Co-immunoprecipitation (Co-IP) and Protein ligation assay (PLA). Empty vector and Flag-AnxA6 transfected MDA-468 cells were grown to 70% confluency and either treated with or without DSP. Immunoprecipitation of AnxA6 from cell lysates was carried out by using anti-flag M2 antibody or anti-AnxA6 antibody **(A)**. Detection of AnxA6 and SNAP23 in the immune complexes was carried out by western blotting. **(B)** Proximity ligation assay (PLA) confirmed the interaction of AnxA6 with SNAP23. All experiments were repeated three times, and representative results are presented. NXL: Non-cross linked, XL: Cross-linked. **(C)** Validation of the interaction of AnxA6 with SNAP23 by GST Pulldown Assay. GST and GST-AnxA6 were expressed in E. coli BL21 and purified to homogeneity. Equal amounts of purified GST and GST-AnxA6 were used to pull down assays using cell lysates from MDA-468. Ponceau stained membrane. **(D)** Membranes were probed with antibodies against the indicated proteins.

**Figure 5.**
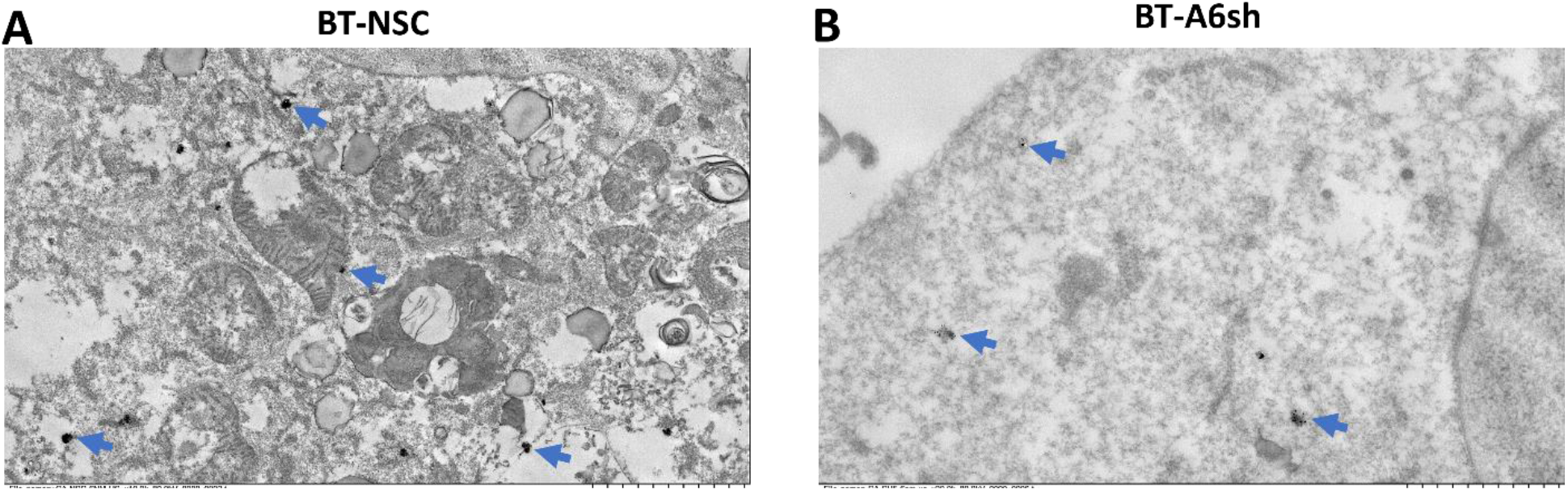
Ca^2+^-dependent localization of AnxA6 on membrane-bound cellular structures. Control AnxA6 expressing and AnxA6 depleted BT-549 cells were treated with (**A** and **B**) 5 mM Ca^2+^ for 5 min, fixed, and processed for immunogold TEM. Images are representative transmission electron micrographs at the same magnification. Arrows indicate the location of AnxA6 within each cell.

### 3.3 AnxA6 neutralizing antibodies strongly inhibit TNBC cell viability

To demonstrate the potential for extracellular AnxA6 to influence the viability of TNBC cells, we assessed the effect of a mouse monoclonal anti-Annexin VI Antibody IgG_2b_ κ (G-10) as neutralizing AnxA6 antibodies (Santa Cruz Biotechnology Inc. SC-166807). We tested the antibody on BT-549 cells that express relatively high levels of AnxA6 and MDA-468 cells that express relatively lower levels of AnxA6. We first show that the antibody recognized AnxA6 on the surface of BT-549 cells (**Fig. 6A**), and then showed that although anti-AnxA6 antibodies dose dependently reduced the viability of TNBC cells, the mesenchymal-like HCC70 and BT-549 cells were more resistant than the epithelial MDA-468 cells (**Fig. 6B**). We next compared the effect of the neutralizing antibodies to the isotype control and show that the viability of BT-549 cells was slightly decreased by 10 μg/ml (**Fig. 6C**), while that of MDA-468 was significantly reduced by 2 μg/ml (range 2-6 μg/ml (**Fig. 6D**). Together this suggests that extracellular AnxA6 is important in the survival of basal-like TNBC cells

## Discussion

Thus far, substantial evidence supports the role of Annexins including AnxA6 in the formation of membrane contact sites (MCS), intraluminal vesicles (ILV), and the transport of cholesterol from late endosomes (23) Several studies have, also reported the potential for AnxA6 enriched exosomes/EVs to influence the progression, drug resistance, and metastasis of breast, pancreatic and other cancers (22, 20, 21, 1). Our study therefore demonstrates that the secretion of EVs and proinflammatory cytokines such as MCP-1, as well as cholesterol enrichment in EVs are AnxA6 dependent. Downregulation of AnxA6 inhibited, while overexpression or chronic lapatinib-induced expression of AnxA6 promoted the secretion of MCP-1, and enrichment of cholesterol in EVs. This study also shows that AnxA6 influences the secretion of these factors by its Ca^2+^-dependent translocation to the plasma membrane and several membrane-bound organelles and that its interaction with SNAP23, a critical component of the SNARE membrane fusion machinery (27) supports at least in part its involvement in the secretion of these factors. Given that AnxA6 is also detected extracellularly and is associated with EVs (7, 8, 9) decrease in cell survival following treatment of TNBC cells with neutralizing anti-AnxA6 antibodies supports the notion that extracellular and/or AnxA6 associated with EVs could be targeted to attenuate TNBC progression, metastasis, and drug resistance.

**Figure 6.**
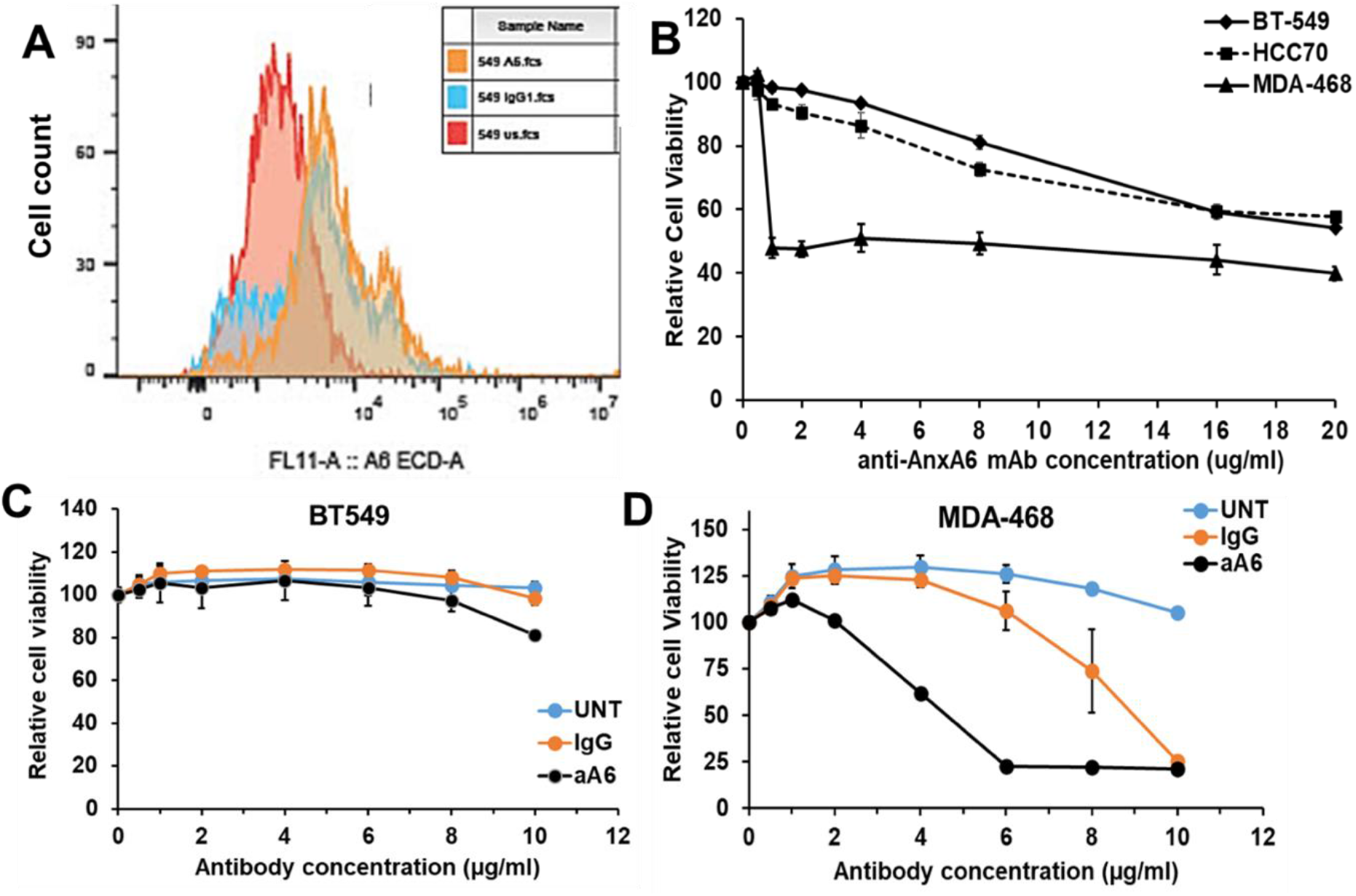
AnxA6 neutralizing antibodies strongly reduce TNBC cell viability. Cell count in various samples **(A)**. Cell viability is reduced in MDA-468 compared to HCC70 and BT-549 **(B)**. Cell viability is lower in AnxA6 neutralizing antibody treated BT-549 compared to untreated and IgG BT-549 cells **(C)**. In MDA-468, AnxA6 neutralizing antibody strongly inhibited cell viability when compared to the untreated and IgG **(D)**. * denotes p<0.05; ** denotes p<0.01; *** denotes p<0.001.

The requirement of Ca^2+^ in AnxA6 mediated exocytosis/secretion of EVs has thus far been shown to depend on the cell type and/or pool of secretory molecules. In HCT116 colon cancer cells, AnxA6 is recruited to MVBs in the presence of Ca^2+^ and that AnxA6 is required for Ca^2+^-dependent exosome secretion both in intact and in permeabilized cells (24). This study specifically demonstrated the requirement of AnxA6 in plasma membrane damage repair-induced secretion of exosomes (24). On the contrary, other reports have suggested that membrane proximal lysosomes rather than multivesicular bodies (MVBs) are responsible for Ca^2+^ dependent exocytosis in non-stimulatory cells (27, 28). Despite these contradictions, rapid release of EVs has been linked to elevated levels of cytosolic Ca^2+^ in T cells and leukemia cells (30, 31). Our immunogold transmission electron micrographs support the notion that in AnxA6 expressing cells, extracellular Ca^2+^ mediated increase in cytosolic Ca^2+^ triggered the translocation of AnxA6 to the plasma membrane and several membrane bound compartments. This effect was markedly reduced following AnxA6 downregulation and associated with decreased secretion of both EVs and proinflammatory cytokines.

Earlier studies reported two modes of Ca^2+^ dependent association of AnxA6 with cell membranes and suggested the involvement of AnxA6 in secretion/exocytosis. At low Ca^2+^-concentrations (up to 150 μM), AnxA6 is thought to bind to the same membrane via its two coplanar annexin modules, but at high Ca^2+^-concentrations (≥ 2 mM), AnxA6 binds two adjacent phospholipid membranes with positive cooperativity (32, 33). However, the argument for negative regulation of secretion by AnxA6 is based on the observation that AnxA6 more effectively inhibited synexin (AnxA7) induced granule aggregation in chromaffin cells. In another study in PC12 cells, overexpression of AnxA6 isoforms almost completely inhibited, while AnxA6 knock-down in these cells was accompanied by a 20% enhancement of dopamine secretion (34). In Chinese hamster ovary cells expression of AnxA6 led to sequestration of cholesterol in the late endosomal compartment and reduced cholesterol in the plasma membrane that negatively affected secretion (35). These reports together, emphasize the notion that subcellular distribution of cholesterol is strongly influenced by the expression levels and subcellular localization of AnxA6 (36, 37).

Thus far, the involvement AnxA6 in the secretion of cytokines is poorly understood and limited to IL-2 secretion studies in Jurkat T cells. In these cells, the intracellular localization of AnxA6 switched from the cytosol to vesicular structures near the plasma membrane following an increase in intracellular Ca^2+^ or decrease in acidity, suggesting that AnxA6 promotes Ca^2+^-and proton-dependent secretion of cytokines (38). This is supported by the observation that T cells isolated from Anxa6 deficient mice exhibited reduced membrane order, low cholesterol levels and reduced interleukin-2 (IL-2) signaling compared with T cells derived from wild type mice (39). Similar observations have been reported following cholesterol depletion with methyl-β-cyclodextrin and statins that led to decreased chemotactic response of monocytes to MCP-1 (40), and that cholesterol rich oxidized low density lipoproteins (OxLDL)-bound MCP-1 retains its ability to recruit monocytes (41). While this supports the role of cholesterol in MCP-1 secretion and chemotactic functions, our finding that the secretion of MCP-1 as well as AnxA6 and cholesterol enriched EVs is AnxA6 dependent further suggests that AnxA6 and cholesterol together influence the secretion and activity of cytokines.

The mechanism by which AnxA6 influences the secretion of cytokines and/or EVs remains poorly understood. Previous studies reported the mislocalization of components of the SNARE (soluble N-ethylmaleimide-sensitive fusion protein attachment protein receptors) complex including SNAP23, STX4, VAMP8 following accumulation of cholesterol in late endosomes (42). Other studies have suggested the regulation of the SNARE complex by several factors denoted as tethering factors that facilitate the fusion of vesicular and plasma membranes (43). The interaction of AnxA6 with SNAP23 as demonstrated in this study links AnxA6 to the SNARE complex, the major membrane fusion machinery that is critical for secretion/exocytosis (44). Given the potential role that Ca^2+^ plays in the fusion of MVBs to the plasma membrane for exocytosis (30) and the Ca^2+^-dependent translocation of AnxA6 to cellular membranes (9), it is possible that the interaction of AnxA6 with SNAP23 either stabilizes the SNARE complex on the plasma membrane, or Ca^2+^-dependently associates with both plasma and vesicular membranes to facilitate vesicle or MVB tethering for effective secretion or exocytosis. However, further studies are necessary to unequivocally demonstrate how AnxA6 influences SNAP23 in the SNARE complex assembly and function in the release of cytokines and/or EVs.

AnxA6 is a predominantly intracellular scaffolding protein with no enzymatic activity that has been shown to influence tumor growth, metastasis and drug resistance (4), but therapeutic targeting of this scaffolding protein has not been amply explored. Our findings that AnxA6 is associated with EVs and previous studies showing that AnxA6 is also a component of EVs and detected in extracellular media EVs (7, 8, 9) suggest that AnxA6 exhibits extracellular functions in TNBC cells. The decrease in the survival of AnxA6 low TNBC cells but not tumor cells expressing relatively high levels of AnxA6 in the presence of monoclonal anti-AnxA6 neutralizing antibodies suggest the requirement for extracellular AnxA6 in the growth of these tumor cells. This is consistent with previous studies in which anti-AnxA2 antibodies were demonstrated to significantly decrease invasion of chick embryo chorioallantoic membrane, as well as the tumor growth and metastasis in breast, ovarian and pancreatic cancers (45, 46, 47). While some of these effects were mediated via annexins as membrane repair proteins (24, 48), our study shows that reducing the levels of extracellular AnxA6 with anti-AnxA6 neutralizing antibodies is a viable therapeutic strategy to target this scaffolding protein especially in the AnxA6-low basal-like breast cancer.

In summary, our data support the model that AnxA6 Ca^2+^-dependently translocates to fusion competent membrane microdomains where it interacts with SNAP23 to stabilize the SNARE complex and/or help in the tethering of vesicles/MVBs to the plasma membrane for exocytosis. Further studies are necessary to determine if AnxA6 is necessary for regulated or constitutive secretion of exosomes and/or proinflammatory cytokines.

## Author contributions

Conceptualization, A.M.S.

Methodology, N.I.S., N.B.V., P.J.B., D.D.B., P.T., H.K.B., J.O., A.J.H., and A.M.S.

Validation, N.I.S., P.T., and A.M.S. Formal Analysis, N.I.S., and A.M.S.

Investigation, N.I.S., P.T., H.K.B., and A.M.S.

Wrote the paper–original draft, N.I.S. and A.M.S.; Review & editing, N.I.S., A.J.H., and A.M.S. Funding acquisition, A.M.S.

Supervision, J.O. and A.M.S.

All authors have read and agreed to the published version of the manuscript.

## References

1. Cao J, Wan S, Chen S, Yang L. ANXA6: a key molecular player in cancer progression and drug resistance. Discov Oncol. 2023;14(1):53.

2. Korolkova OY, Widatalla SE, Whalen DS, Nangami GN, Abimbola A, Williams SD, et al. Reciprocal expression of Annexin A6 and RasGRF2 discriminates rapidly growing from invasive triple negative breast cancer subsets. PLoS One. 2020;15(4):e0231711.

3. Whalen DS, Widatalla SE, Korolkova OY, Nangami GS, Beasley HK, Williams SD, et al. Implication of calcium activated RasGRF2 in Annexin A6-mediated breast tumor cell growth and motility. Oncotarget. 2019;10(2):133–51.

4. Korolkova OY, Widatalla SE, Williams SD, Whalen DS, Beasley HK, Ochieng J, et al. Diverse Roles of Annexin A6 in Triple-Negative Breast Cancer Diagnosis, Prognosis and EGFR-Targeted Therapies. Cells. 2020;9(8).

5. Widatalla SE, Korolkova OY, Whalen DS, Goodwin JS, Williams KP, Ochieng J, et al. Lapatinib-induced annexin A6 upregulation as an adaptive response of triple-negative breast cancer cells to EGFR tyrosine kinase inhibitors. Carcinogenesis. 2019;40(8):998–1009.

6. Williams SD, Smith TM, Stewart LV, Sakwe AM. Hypoxia-Inducible Expression of Annexin A6 Enhances the Resistance of Triple-Negative Breast Cancer Cells to EGFR and AR Antagonists. Cells. 2022;11(19).

7. Ochieng J, Pratap S, Khatua AK, Sakwe AM. Anchorage-independent growth of breast carcinoma cells is mediated by serum exosomes. Exp Cell Res. 2009;315(11):1875–88.

8. Koumangoye RB, Nangami GN, Thompson PD, Agboto VK, Ochieng J, Sakwe AM. Reduced annexin A6 expression promotes the degradation of activated epidermal growth factor receptor and sensitizes invasive breast cancer cells to EGFR-targeted tyrosine kinase inhibitors. Mol Cancer. 2013;12(1):167.

9. Sakwe AM, Koumangoye R, Guillory B, Ochieng J. Annexin A6 contributes to the invasiveness of breast carcinoma cells by influencing the organization and localization of functional focal adhesions. Exp Cell Res. 2011;317(6):823–37.

10. Singh S, Anshita D, Ravichandiran V. MCP-1: Function, regulation, and involvement in disease. Int Immunopharmacol. 2021;101(Pt B):107598.

11. Carr MW, Roth SJ, Luther E, Rose SS, Springer TA. Monocyte chemoattractant protein 1 acts as a T-lymphocyte chemoattractant. Proc Natl Acad Sci U S A. 1994;91(9):3652–6.

12. Xu LL, Warren MK, Rose WL, Gong W, Wang JM. Human recombinant monocyte chemotactic protein and other C-C chemokines bind and induce directional migration of dendritic cells in vitro. J Leukoc Biol. 1996;60(3):365–71.

13. Fousek K, Horn LA, Palena C. Interleukin-8: A chemokine at the intersection of cancer plasticity, angiogenesis, and immune suppression. Pharmacol Ther. 2021;219:107692.

14. Shen S, Song Y, Zhao B, Xu Y, Ren X, Zhou Y, et al. Cancer-derived exosomal miR-7641 promotes breast cancer progression and metastasis. Cell Commun Signal. 2021;19(1):20.

15. O’Byrne KJ, Dalgleish AG, Browning MJ, Steward WP, Harris AL. The relationship between angiogenesis and the immune response in carcinogenesis and the progression of malignant disease. Eur J Cancer. 2000;36(2):151–69.

16. Stow JL, Murray RZ. Intracellular trafficking and secretion of inflammatory cytokines. Cytokine Growth Factor Rev. 2013;24(3):227–39.

17. Lacy P, Stow JL. Cytokine release from innate immune cells: association with diverse membrane trafficking pathways. Blood. 2011;118(1):9–18.

18. Colombo M, Raposo G, Thery C. Biogenesis, secretion, and intercellular interactions of exosomes and other extracellular vesicles. Annu Rev Cell Dev Biol. 2014;30:255–89.

19. Wan S, He QY, Yang Y, Liu F, Zhang X, Guo X, et al. SPARC Stabilizes ApoE to Induce Cholesterol-Dependent Invasion and Sorafenib Resistance in Hepatocellular Carcinoma. Cancer Res. 2024;84(11):1872–88.

20. Leca J, Martinez S, Lac S, Nigri J, Secq V, Rubis M, et al. Cancer-associated fibroblast-derived annexin A6+ extracellular vesicles support pancreatic cancer aggressiveness. J Clin Invest. 2016;126(11):4140–56.

21. Uchihara T, Miyake K, Yonemura A, Komohara Y, Itoyama R, Koiwa M, et al. Extracellular Vesicles from Cancer-Associated Fibroblasts Containing Annexin A6 Induces FAK-YAP Activation by Stabilizing beta1 Integrin, Enhancing Drug Resistance. Cancer Res. 2020;80(16):3222–35.

22. Keklikoglou I, Cianciaruso C, Guc E, Squadrito ML, Spring LM, Tazzyman S, et al. Chemotherapy elicits pro-metastatic extracellular vesicles in breast cancer models. Nat Cell Biol. 2019;21(2):190–202.

23. Rentero C, Blanco-Munoz P, Meneses-Salas E, Grewal T, Enrich C. Annexins-Coordinators of Cholesterol Homeostasis in Endocytic Pathways. Int J Mol Sci. 2018;19(5).

24. Williams JK, Ngo JM, Lehman IM, Schekman R. Annexin A6 mediates calcium-dependent exosome secretion during plasma membrane repair. Elife. 2023;12.

25. Jezequel P, Frenel JS, Campion L, Guerin-Charbonnel C, Gouraud W, Ricolleau G, et al. bc-GenExMiner 3.0: new mining module computes breast cancer gene expression correlation analyses. Database (Oxford). 2013;2013:bas060.

26. Boye TL, Maeda K, Pezeshkian W, Sonder SL, Haeger SC, Gerke V, et al. Annexin A4 and A6 induce membrane curvature and constriction during cell membrane repair. Nat Commun. 2017;8(1):1623.

27. Faigle W, Colucci-Guyon E, Louvard D, Amigorena S, Galli T. Vimentin filaments in fibroblasts are a reservoir for SNAP23, a component of the membrane fusion machinery. Mol Biol Cell. 2000;11(10):3485–94.

28. Jaiswal JK, Chakrabarti S, Andrews NW, Simon SM. Synaptotagmin VII restricts fusion pore expansion during lysosomal exocytosis. PLoS Biol. 2004;2(8):E233.

29. Jaiswal JK, Andrews NW, Simon SM. Membrane proximal lysosomes are the major vesicles responsible for calcium-dependent exocytosis in nonsecretory cells. J Cell Biol. 2002;159(4):625–35.

30. Messenger SW, Woo SS, Sun Z, Martin TFJ. A Ca(2+)-stimulated exosome release pathway in cancer cells is regulated by Munc13-4. J Cell Biol. 2018;217(8):2877–90.

31. Savina A, Furlan M, Vidal M, Colombo MI. Exosome release is regulated by a calcium-dependent mechanism in K562 cells. J Biol Chem. 2003;278(22):20083–90.

32. Buzhynskyy N, Golczak M, Lai-Kee-Him J, Lambert O, Tessier B, Gounou C, et al. Annexin-A6 presents two modes of association with phospholipid membranes. A combined QCM-D, AFM and cryo-TEM study. J Struct Biol. 2009;168(1):107–16.

33. Zaks WJ, Creutz CE. Annexin-chromaffin granule membrane interactions: a comparative study of synexin, p32 and p67. Biochim Biophys Acta. 1990;1029(1):149–60.

34. Podszywalow-Bartnicka P, Kosiorek M, Piwocka K, Sikora E, Zablocki K, Pikula S. Role of annexin A6 isoforms in catecholamine secretion by PC12 cells: distinct influence on calcium response. J Cell Biochem. 2010;111(1):168–78.

35. Cubells L, Vila de Muga S, Tebar F, Wood P, Evans R, Ingelmo-Torres M, et al. Annexin A6-induced alterations in cholesterol transport and caveolin export from the Golgi complex. Traffic. 2007;8(11):1568–89.

36. Adjibade P, Simoneau B, Ledoux N, Gauthier WN, Nkurunziza M, Khandjian EW, et al. Treatment of cancer cells with Lapatinib negatively regulates general translation and induces stress granules formation. PLoS One. 2020;15(5):e0231894.

37. Emeriau N, de Clippele M, Gailly P, Tajeddine N. Store operated calcium entry is altered by the inhibition of receptors tyrosine kinase. Oncotarget. 2018;9(22):16059–73.

38. Podszywalow-Bartnicka P, Strzelecka-Kiliszek A, Bandorowicz-Pikula J, Pikula S. Calcium- and proton-dependent relocation of annexin A6 in Jurkat T cells stimulated for interleukin-2 secretion. Acta Biochim Pol. 2007;54(2):261–71.

39. Cornely R, Pollock AH, Rentero C, Norris SE, Alvarez-Guaita A, Grewal T, et al. Annexin A6 regulates interleukin-2-mediated T-cell proliferation. Immunol Cell Biol. 2016 Jul;94(6):543–53.

40. Saha AK, Mousavi M, Dallo SF, Evani SJ, Ramasubramanian AK. Influence of membrane cholesterol on monocyte chemotaxis. Cell Immunol. 2018 Feb;324:74–77.

41. Wiesner P, Tafelmeier M, Chittka D, Choi SH, Zhang L, Byun YS, et al. MCP-1 binds to oxidized LDL and is carried by lipoprotein(a) in human plasma. J Lipid Res. 2013 Jul;54(7):1877–83.

42. Enrich C, Rentero C, Hierro A, Grewal T. Role of cholesterol in SNARE-mediated trafficking on intracellular membranes. J Cell Sci. 2015;128(6):1071–81.

43. Ungermann C, Kummel D. Structure of membrane tethers and their role in fusion. Traffic. 2019;20(7):479–90.

44. Jahn R, Cafiso DC, Tamm LK. Mechanisms of SNARE proteins in membrane fusion. Nat Rev Mol Cell Biol. 2024 Feb;25(2):101–118.

45. Gounou C, Rouyer L, Siegfried G, Harté E, Bouvet F, d’Agata L, et al. Inhibition of the membrane repair protein annexin-A2 prevents tumor invasion and metastasis. Cell Mol Life Sci. 2023 Dec 13;81(1):7.

46. Lokman NA, Elder AS, Ween MP, Pyragius CE, Hoffmann P, Oehler MK, et al. Annexin A2 is regulated by ovarian cancer-peritoneal cell interactions and promotes metastasis. Oncotarget. 2013 Aug;4(8):1199–211.

47. Zheng L, Foley K, Huang L, Leubner A, Mo G, Olino K, et al. Tyrosine 23 phosphorylation-dependent cell-surface localization of annexin A2 is required for invasion and metastases of pancreatic cancer. PLoS One. 2011 Apr 29;6(4):e19390.

48. Demonbreun AR, Fallon KS, Oosterbaan CC, Bogdanovic E, Warner JL, Sell JJ, et al. Recombinant annexin A6 promotes membrane repair and protects against muscle injury. J Clin Invest. 2019 Nov 1; 129(11):4657–4670.

